# Distinction between slow waves and delta waves sheds light to sleep homeostasis and their association to hippocampal sharp waves ripples

**DOI:** 10.1101/2022.12.27.522034

**Authors:** Karim El-Kanbi, Gaëtan de Lavilléon, Sophie Bagur, Marie Lacroix, Karim Benchenane

## Abstract

Cortical slow waves and delta waves are hallmarks of NREM sleep and key elements in physiological processes such as memory consolidation and sleep homeostasis. However, no clear distinction has been made between the two types of electrophysiological events. We introduce a new distinction between slow waves, large amplitude waves on filtered LFP cortical signals, and delta waves, characterized by an inversion between deep and superficial layers and the co-occurrence with a cortical down state. Detection of slow waves, as large amplitude waves during NREM sleep, reveals that a large portion of them does not share the properties of delta waves and does not co-occur with cortical down states. Therefore, we distinguish type-1 slow waves, that are associated to a down state, from type-2 slow waves. We show that 1) only type-1 slow waves undergo strong homeostatic regulations and 2) type-2 slow waves create confusion about the temporal link with hippocampal sharp-wave ripples. Thus, we confirm that delta waves detected with our new method can be used as a proxy for down state. Altogether, this confirms the importance of a robust and accurate definition of delta waves to understand the fine neurophysiological mechanisms at stake during NREM sleep.

## Introduction

Since the earliest investigations of brain electrical activity, researchers have noticed that changes during the transition to sleep as well as changes in consciousness level are associated with modified EEG patterns, both in animal models [1] and in humans [2].

In humans, NREM sleep is characterized by the occurrence of spindles, K-complexes and slow oscillations. As the sleep state deepens, the number of K-complexes/delta waves progressively increases and gives rise to slow oscillations. Thus, the deepest stage of sleep is called slow wave sleep (SWS). More precisely, SWS in humans is defined by the presence of high-voltage (> 75 μV) synchronized EEG waveforms: delta oscillations (1-4 Hz) and slow oscillations (SO) (< 1Hz), for which the power densities in the 0.75-4 Hz EEG range are typically referred to as slow wave activity (SWA). SWS is thus defined by electrophysiological characteristics that can be visually assessed and scored based on the frequency and the amplitude of brain oscillations, according to standardized criteria [3].

Aside from some minor differences, brain oscillations and the electrophysiological events observed in humans are extremely well conserved in mammals, especially rodents [4]. In both humans and rodents, sleep is organized into cycles of alternating REM and NREM sleep. The cycle duration is around 90 min in humans and 10 min in rodents [5]. The most striking difference between human and rodent sleep lies in the overall architecture: while sleep occurs in generally one monophasic block during the night in humans, the primarily nocturnal rodents display a highly fragmented polyphasic sleep, which can occur both during the day and the night. In addition, NREM sleep sub-stages appear to be much more blurred in rodents [6]. This explains why many authors use the term SWS in rodents to describe the whole NREM sleep [6].

### Role of delta waves and slow oscillations on memory consolidation during sleep

Different parameters that characterize SWS are been shown to be involved in memory consolidation. Indeed, the slow oscillation (defined as the spectral power <1 Hz) following intense declarative learning is increased [7]; furthermore, the increase in coherence is mostly observed at a phase of the SO corresponding to cortical up-states, an effect almost identical in humans and rodents [8]. A separate study has found that the length of the up-state estimated by the positive wave of the slow oscillations directly relates to overnight memory improvements in a declarative memory task [9]. Moreover, in an implicit motor task, during which subjects implicitly learned to adapt their movements to a rotated display, there is a local increase in SWA during sleep in the very same brain areas that had previously been activated during the task [10]. Importantly, the improved performance in the rotation adaptation task after sleep was predicted by the increase in SWA. Furthermore, one study showed that enhancing the slow oscillation with a close-loop device can improve the beneficial effect of sleep on declarative memory. Stimuli were applied in a time-locked manner to the SO up-phase [11]. The enhancement of the SO was associated with an increase in spindles. However, it is important to note that it is still debated since other studies failed to find the same effect [12].

The main hypothesis, regarding the influence of SO on memory processes, is that slow cortical rhythms help for the global coordination of thalamocortical rhythms and hippocampal events [13], [14]. This favors the interaction between the hippocampus and the cortex, that has been shown to be crucial for memory consolidation. Indeed it has been shown in rodent that wake experiences are replayed during hippocampal sharp waves ripples (SWR) and that these replays are associated with reactivation in the prefrontal cortex [15]. The coordination between the cortical rhythms and the hippocampal SWR is thus crucial. Accordingly, enhancing the coordination between the SWR and cortical delta waves in rodents enhances the beneficial effect of sleep on memory [16]. However, even if this hypothesis is attractive, there are still some weaknesses and unresolved issues. First of all, the coordination between these rhythms is extremely week. Less than 5% of ripples are found to be in the SWR-delta wave-spindles triplets [17]. Moreover, some studies found that hippocampal SWRs occur before the delta wave/up states [14], [17], some others found the exact opposite [18], [19]. The interpretation of these conflicting results is made difficult by the different methods of detection and terminology used in the different studies.

### Role of delta/waves/K complex and slow oscillations on sleep homeostasis

Even if slow cortical rhythms have been involved in memory processes, their strongest characteristic is their very tight link with sleep homeostasis. The amount of slow waves is strongly increased at the beginning of sleep and decreases progressively during sleep [20], [21]. Moreover, longer periods of wakefulness before sleep are associated with a higher density of slow waves at the beginning of sleep. It has thus been proposed that slow waves are a marker of sleep pressure and sleep need [20]. Computational models have shown that this can be modeled with exponential fits, and the density of slow waves is considered to represent the process S that is an indicator of sleep pressure [22], [23]. From the observation of their occurrence, slow waves seem to correspond to what is excepted from a process that cures the brain of the lack of sleep. However, on the contrary to their role in memory processes, there is still no hypothesis on the role of slow waves on the restorative function of sleep.

### Towards a better definition of NREM sleep rhythms in human and rodent

One of the main pitfalls when trying to understand the precise role and the action mechanism of slow rhythms is their lack of precise definition. Indeed, even if brain oscillations are very similar in most mammalian species, including the various rhythms observed during SWS [4], the precise definition of each of the EEG events is still rather vague. Accordingly, it is sometimes difficult to define precisely the differences between slow waves, slow oscillation, delta rhythm, K-complexes or delta waves. Perhaps the electrophysiological event that has been defined with the highest precision is the delta wave. Delta waves correspond to a sharp deflection of the LFP that lasts around (200-500ms) corresponding to the classical range for delta (2-4 Hz) [24]. Interestingly, while most sleep studies in rodents use the term ‘delta wave’, very few have used the term ‘K-complex’ which is mostly used in human studies. One study specifically investigated K-complex waveforms based on the shape of the wave [25], although this report stands out as an exception in the rodent literature. Concerning the slow oscillation, its frequency defined as the lowest peak in the power spectrum is slightly faster in rodents than in humans (1.35 Hz vs. 0.8Hz) [8].

### Neurophysiological mechanisms associated with delta waves and slow oscillations

The seminal work of Mircea Steriade showed that slow oscillations are based on alternating up- and down-states of the membrane potential of cortical neurons that are not seen during waking or REM sleep [26], [27]. By looking more closely, it appears that this alternation occurs at the time of delta waves, and recording a large population of neurons showed that this is associated with an OFF period with all cortical neurons being silent. The slow oscillation would thus correspond to the underlying process that controls the occurrence of delta waves [28].

Recordings at different depths of the cortex showed that down states/Off periods are detectable in local field potentials (LFP)/EEG as a positive wave in the deep layers of the cortex and as a negative wave in the surface layers, which looks like an electrical dipole between the two recordings sites [29]. These LFP events are then called delta waves. It is still unclear how a cessation of neuronal activity in the cortex could be linked to a dipole that is visible in the LFP signal [30]. Nevertheless, this implies that recording at different cortical depths can disambiguate the delta waves from random fluctuations.

However, this raises the question of whether all slow waves correspond to delta waves or not. We will attempt to provide a better characterization of the fluctuations of large amplitude and slow frequency that occurs during the NREM sleep and whether the occurrence of the different types of waves are associated with an electrical dipole between the superficial and the deep layers, or with the ON/OFF fluctuations in the cortical neuronal activity. We will then investigate the relationship of the different types of slow waves with sleep homeostasis and with the hippocampal SWRs.

## Results

NREM sleep is associated with low frequency and large amplitude fluctuations visible in the LFP or EEG. To characterize these fluctuations, we recorded ECoG in the frontal cortex, LFP in the superficial and deep layers of the prefrontal cortex (PFC) and large ensembles of neurons with tetrodes in the PFC.

We noticed that the fluctuations observed during NREM sleep can be classified into two categories of waves. The first one corresponds to the typical delta waves with a negative deflection in the superficial layers and positive deflections in the deep layers. They are associated with a silence of the neuronal population in the PFC, corresponding to the OFF periods and likely associated with down states. The second category corresponds to the slow wave, which is a large amplitude wave recorded in a single location. However, a large amount of large amplitude fluctuations does not match this criterion and show no dipole and no obvious modification of the neuronal activity. This would thus correspond to slow waves but not with delta waves (Figure 1). In the manuscript, we will distinguish two types of slow waves: type-1 slow wave which corresponds to the co-occurrence of a slow wave and a down state, and type-2 slow wave which refers to the detection of a slow wave without the observation of a down state. Importantly, most of the methods used to detect slow waves are based on a threshold applied to the filtered signal in the low frequency band and thus will fail to dissociate the two categories of slow waves.

**Figure 1:**
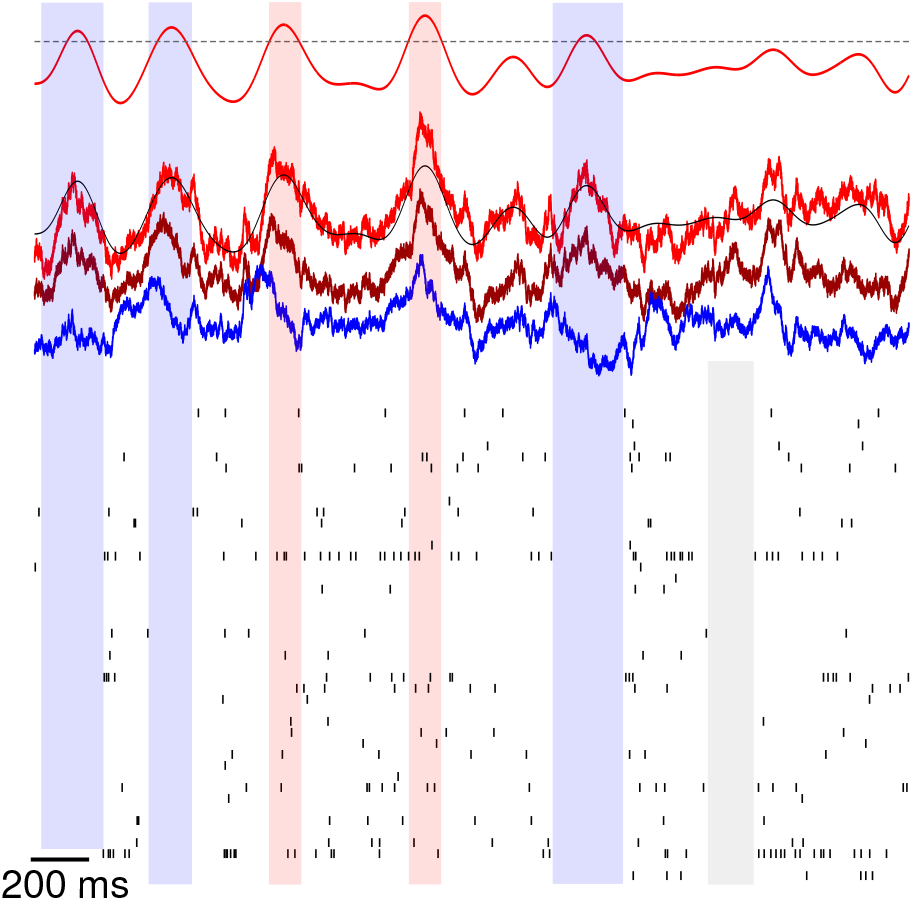
Traces of neocortical recordings during NREM sleep. (top) Filtered LFP signal and threshold for SW detection, (middle) LFP traces from different depths, (bottom) Single-unit activity. Blue rectangles indicate the simultaneous detection of a down state and a slow wave (type-1 slow wave), red rectangles indicate slow waves not associated with a down state (type-2 slow wave), the gray rectangle shows a down state not associated with a slow wave.

### Identification of OFF periods / DOWN states

One of the main characteristics of delta waves is their association with an OFF period. However, a silence of the recorded population of neurons does not necessarily correspond to an OFF period since it strongly depends on the number of neurons recorded. By shuffling the activity of the neurons, OFF periods will be affected only if the number of neurons recorded is large enough to represent the global population (Fig. 2a). We thus characterized the number of neurons required to identify OFF periods with extracellular recordings. We quantified the number of OFF periods of different durations, both during wakefulness and sleep, with and without shuffling. As shown in figure 2b, the distribution of the OFF periods displays a bump for sleep OFF periods, above around 100ms, which is absent during wakefulness. The permutation disturbs the OFF periods selectively for durations above 75ms. Interestingly, both the bump in the distribution and the permutation effect are systematically observed when the number of neurons recorded is above 40. This can be also observed sometimes for a lower number of neurons, but only in specific cases relying on the type of neurons recorded and the total amount of spikes. We, therefore, chose the limit of 75ms to define OFF periods that identify silences of the total population of cortical neurons, that are likely associated with down states of the membrane potential of the cortical neurons. Because of the most common use in the literature, we will use down states in the rest of the manuscript.

**Figure 2-.**
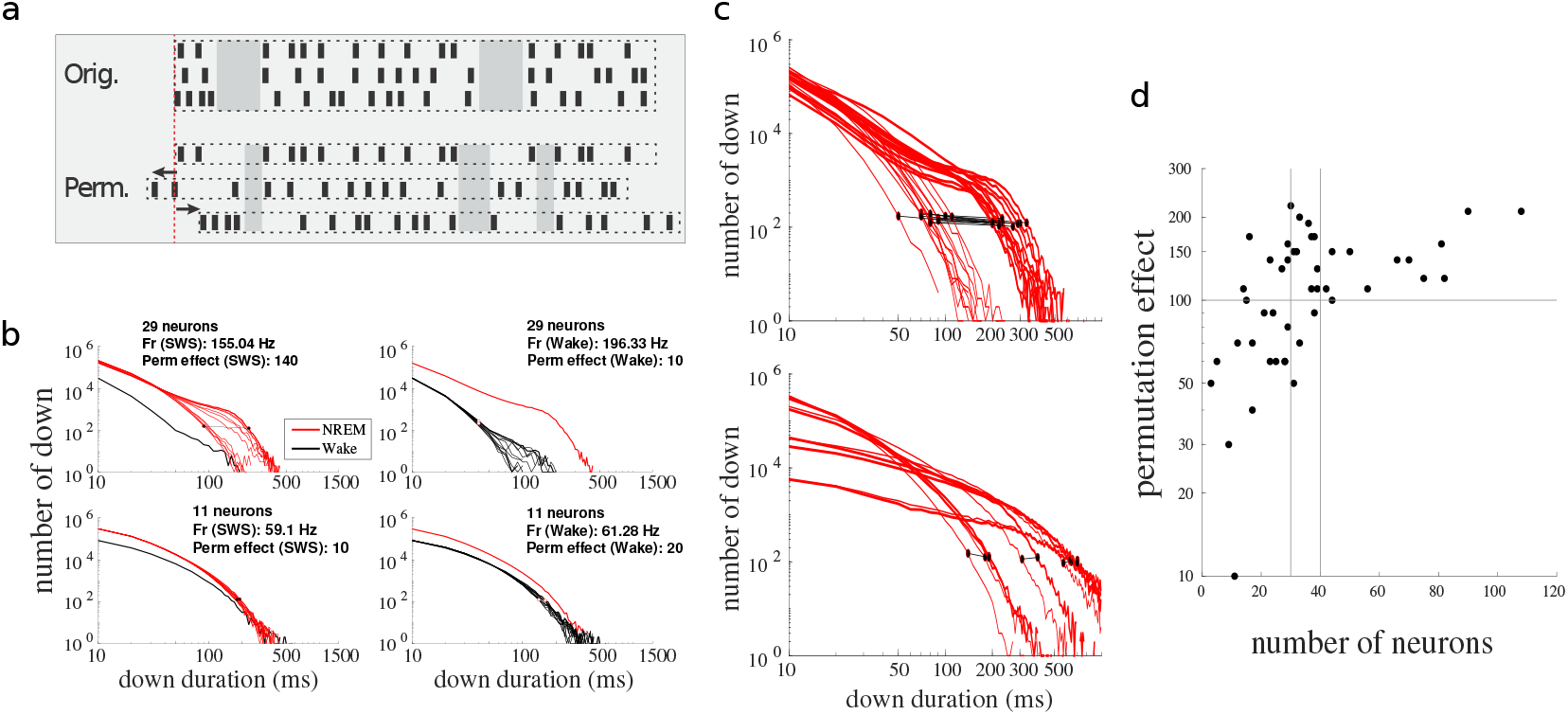
Detecting down states from MUA: minimum firing rate and neurons is required to avoid fake detections. **a**. Schematic of spike trains permutations: as down state is a network phenomenon, permuting each train independently affects distributions of true down states duration. **b**. Histogram of down states duration and effect of permutations. *Top-left*: One record affected by permutations during NREM, *top-right*: permutations during wake on the same record. *Bottom*: One record with only 11 neurons, permutations do not change the distribution of down states durations, which are similar for NREM and Wake. **c**. Effect of permutations on 5 highly permuted records (*top*) and 5 less permuted records (*bottom*). **d**. Permutation effect depends on the number of neurons recorded. Permutation effect corresponds to the distance between the NREM curve and the most permuted curve at y=100. Each point corresponds to a single record (n=19 mice).

**Figure 3-.**
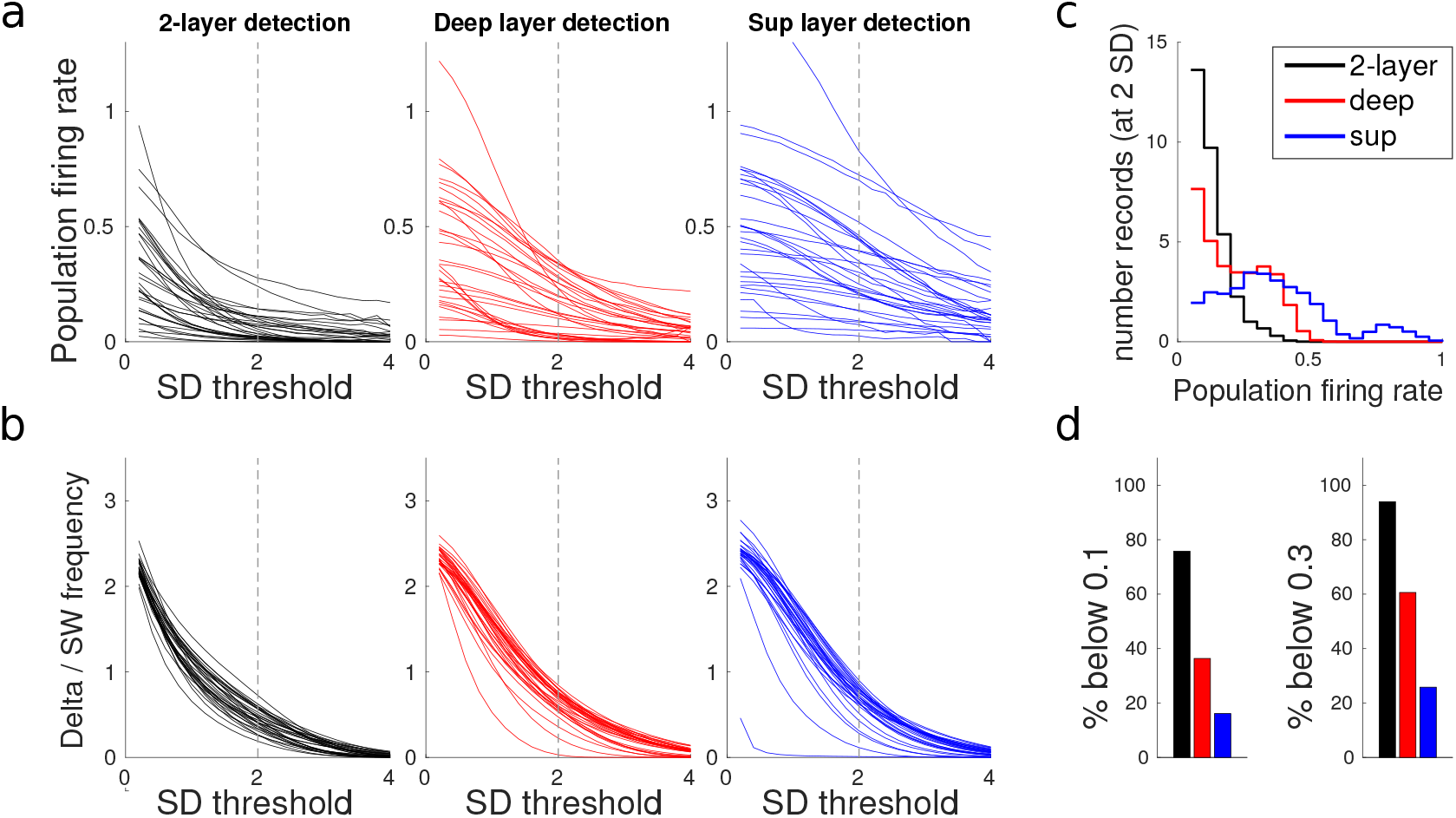
Detection method based on the LFP inversion between deep and superficial layers. *Comparison between the 2-layer method and classical methods on deep or superficial layer*s. **a**. Firing rate on slow / delta waves as a function of the amplitude threshold in SD (standard deviation). The 2-layer detection uses both signals from deep and superficial layers. Each curve corresponds to a sleep session. Firing rate corresponds to the number of spikes per bins of 5ms. **b**. Frequency of event detected (in Hz) for each method of detection. **c**. Histogram of firing rate at 2 SD threshold for the 3 methods. **d**. Percentage of records below 0.1 at 2 SD (left) and below 0.3 (right).

### Identification of delta waves and OFF periods / DOWN states

We then looked for the best method to identify delta waves. We compared waves identified with a single channel, located in superficial or deep layers, and waves detected by using differential recording (2-layers detection). When using a threshold of 2 standard deviations, the 2-layers detection method invariably identifies events associated with a drop of the neuronal activity above 85%. This was rarely observed with recording in the superficial layer, and only for the deepest recordings in the deep layers. We thus used the 2-layer detection method with a threshold of 2 standard deviations for the rest of the study.

### Relationship between the two types of slow waves and the UP/DOWN fluctuations

Most of the studies that investigate the role of NREM sleep on memory or sleep homeostasis focus on large amplitude slow waves without specifying the exact definition of the event considered. Most of the authors considered as a fact that slow waves correspond to Up-Down fluctuation, without providing evidence whether it is the case or not. In other terms, they implicitly admit that all slow waves correspond to delta waves. Yet, the methods used rely on recordings of the EEG, ECoG or single LFP, and are not specific enough to guaranty such a conclusion. For that reason, we used our recordings to provide the recording set-up that ensures the identification of true delta waves associated with down states.

We first identified the down states with the methods described previously. The average at the time of down states of the LFP recorded in different PFC layers leads to events of different amplitudes and different polarities (Fig. 4a). Typically, ECoG and superficial layers display negative deflections of moderate amplitude, while deep layers LFP display positive deflections, with an increasing amplitude with depth. We then defined 6 different groups of recordings based on the amplitude of the signal: ECoG, and groups 1 (superficial) to 5 (deep) (Fig. 4b,c). We then used the classical method used to detect slow waves by filtering the LFP in the low frequency band and using a threshold of 2SD. We observed that only the group 4 and 5 (the deepest recording) are associated with a large decrease in firing rate and show a clear large peak in the cross-correlogram with down states (Fig 4d,e). Quantification showed that most of the situations do not allow a good identification of down states with a false positive rate above 40%. Only the deepest recordings showed a good detection specificity with only 20% (Fig. 4f).

**Figure 4-.**
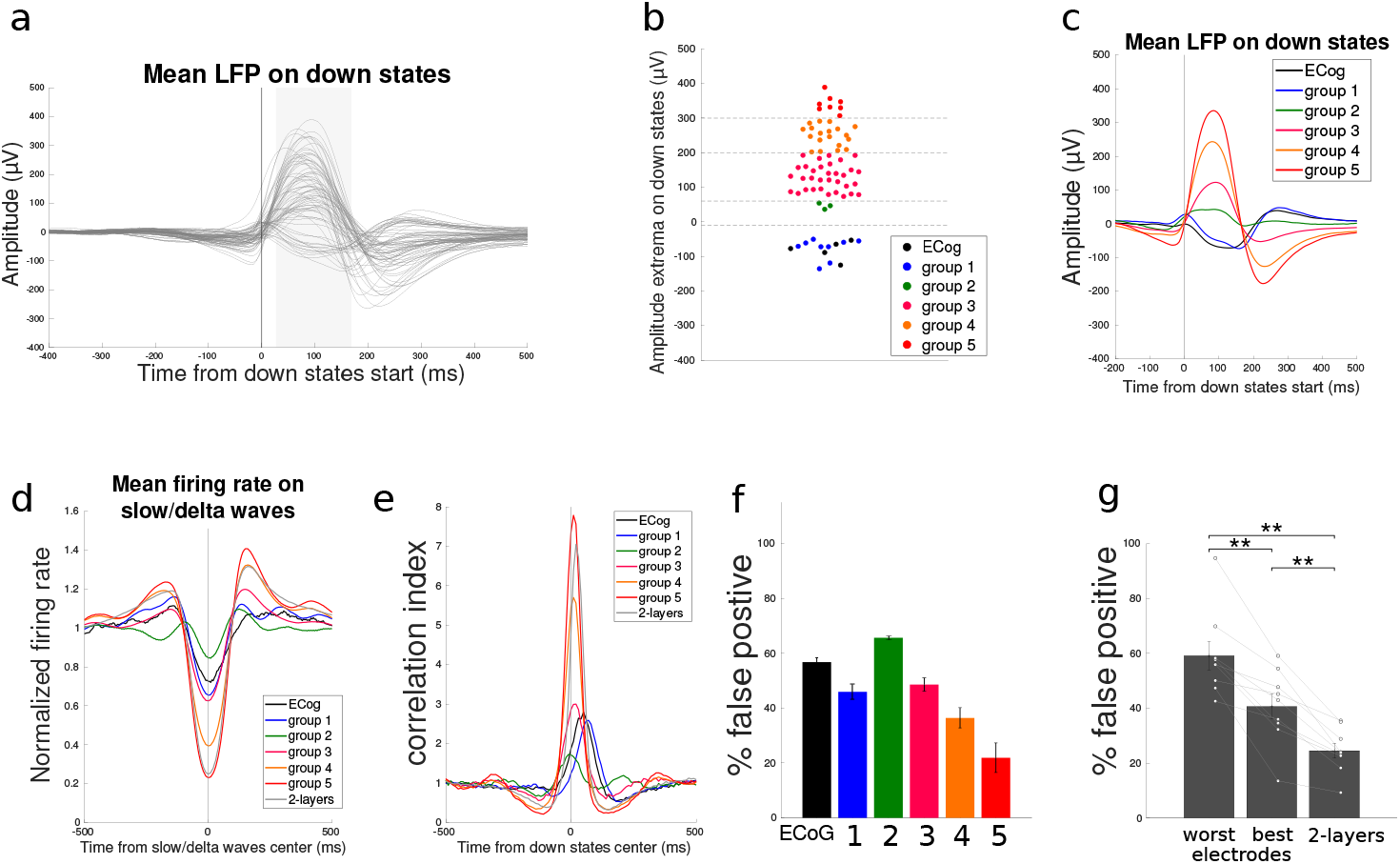
Co-occurrence of slow waves and down states for each group of electrodes - comparison with the 2-layer method. **a**. Mean LFP responses on down state for all the electrodes. **b**. Electrode classification according to the amplitude of extrema in the gray rectangle of graph a. ECog and superficial electrodes (group 1) have negative deflections. **c**. Mean responses on down states for each group. **d**. Normalized firing rate on slow waves detected in each group. 2-layers correspond to delta waves detected with two channels of different depths. **e**. Cross-correlogram between down states and slow waves of each group, comparison with delta waves detected with the 2-layer method. **f**. Percentage of fake detection for each group (Kruskal-Wallis test, p=3,13.10^−5^, Chi^2^=28,33). **g**. Comparison of the percentage of fake detection between the worst electrode, the best electrode and the 2-layers method (Paired Wilcoxon signed-rank test: Signed Rank statistic=45,45,45, p=0.0039,0.0039,0.0039, n=7).

We showed that for each mouse (with only a minority having electrode belonging to the group 5), the 2-layers detection outperforms other methods, having a false detection rate below 20% (Fig. 4g). Importantly, the ECoG, which is close to the EEG classically used in most sleep studies, displays a false detection rate of 60%. In other terms, less than half of the waves detected with EEG correspond to down states (Fig. 4f) and only 44.7 % +/- 2.94 (n=6) correspond to delta waves detected with the 2 layers method.

### Characterization of slow waves: definition of true and false delta waves

Some slow waves do not share the properties of delta waves in terms of inversion between superficial and deep layers or regarding their association with down states. However, it is not clear whether the two types of slow waves have the same physiological functions. We then differentiate two types of slow waves whether they occur with a down state.

First of all, we detected large negative deflections in the superficial layer and considered the 25% waves with the strongest positive (high inversion) and negative (low inversion, or without inversion) deflection in the deep layer (Fig. 5a-b). By using this method, we observed that slow waves with high inversion were associated with a massive decrease in firing rate, a decrease that was absent for the slow waves without inversion (Fig. 5c). Accordingly, 80% of the slow waves with inversion were associated with a down state, while this percentage was only 30% for the slow waves without inversion (Fig. 5d). The same dissociation was done with the large deflections detected in deep layers (Fig. 5e-h) showing similar results. Then, we defined type-1 and type-2 slow waves regarding respectively the presence or the absence of down state at the time of the detected slow wave.

**Figure 5-.**
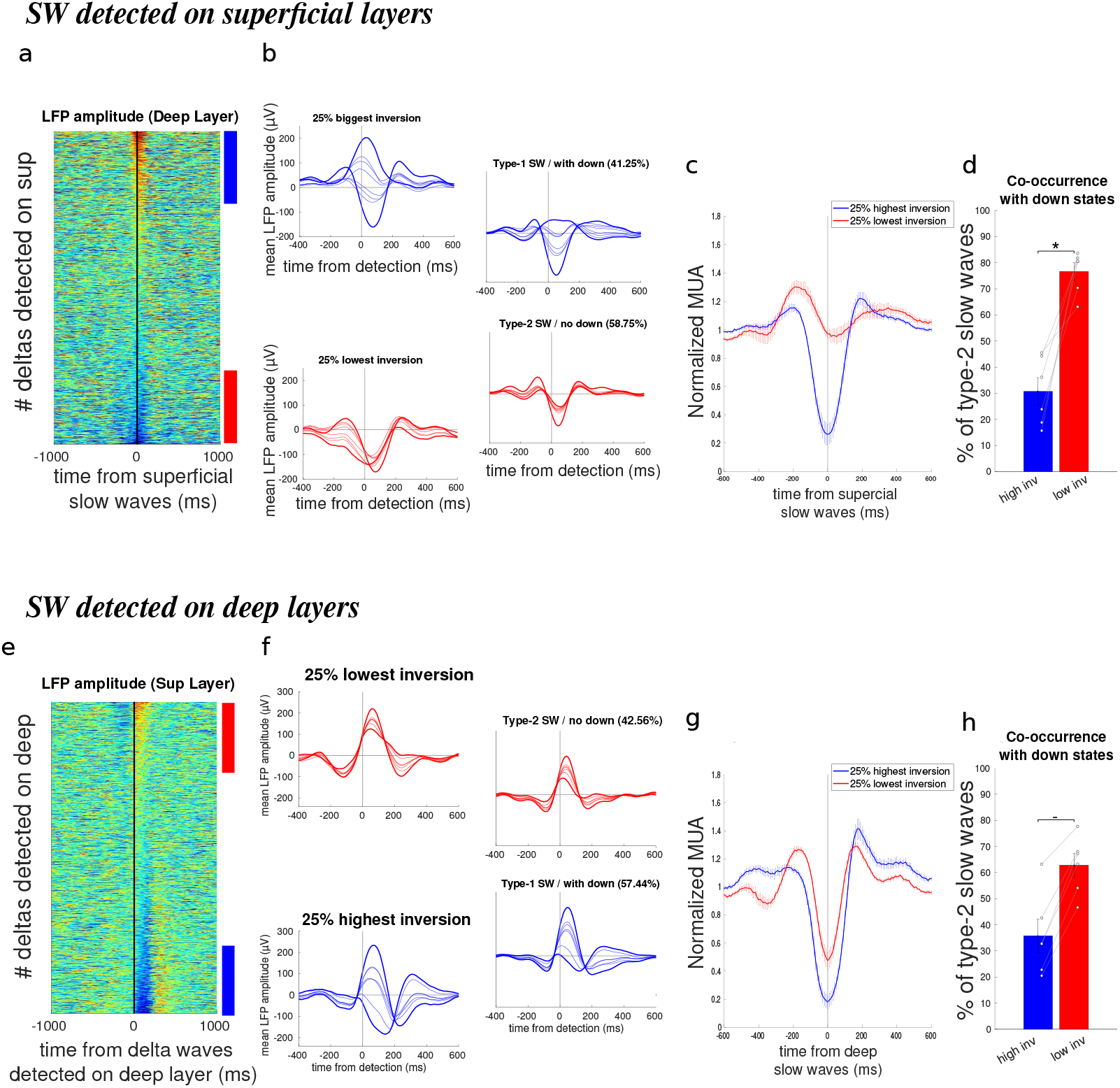
LFP inversion is crucial to detect type-1 slow waves. *(a-d) Slow waves detected in superficial layers*. **a**. Deep layer LFP amplitude for each slow wave detected on the superficial layer, sorted by amplitude in t=0ms (for one sleep session). **b**. Mean LFP amplitude for each channel on slow waves for: 25% highest and lowest inversion (left) and type-1 and type-2 slow waves (right). *Same session as a*. **c**. Normalized firing rate on the highest and lowest inverted slow waves (n=6). **d**. Percentage of slow waves not associated with a down state (Paired Wilcoxon signed-rank test: Signed Rank statistics=0, p=0.031, n=6). *(e-h) Slow waves detected in deep layers*. **e**. Superficial layer LFP amplitude for each slow wave detected on the deep layer, sorted by amplitude in t=50ms (one sleep session). **f**. Mean LFP amplitude for each channel on slow waves for: 25% lowest and highest inversion (left) and type-1 and type-2 slow waves (right). *Same session as f*. **g**. Normalized firing rate on the highest and lowest inverted slow waves (n=6). **h**. Percentage of slow waves not associated with a down state (Paired Wilcoxon signed-rank test: Signed Rank statistics=0, p=0.031, n=6).

### Homeostatic regulation of the two types of slow wave

We then used these definitions of type-1 and type-2 slow waves to investigate their respective homeostatic regulations during sleep. In figure 7a, we display a representative session showing the evolution during sleep of different features: 1) slow wave activity (SWA, defined as in most of the article by the spectral power 0.5-4 Hz), 2) down states, 3) delta waves (defined with the 2 layers method), 4) type-1 slow waves (with down state) and 5) type-2 slow waves (no down state). Three different fits were used: simple linear fit, double linear fit (0-3h and 3h-rest of the sleeping period) and exponential fit. For clarity, the curves displaying the linear fits and the exponential fits are shifted upward and downward. A strong decrease is observed for down states, delta waves and type-1 slow waves while a moderate decrease is observed for the SWA and is almost not visible for type-2 slow waves (Fig. 6a). This difference is observed for all the recording sessions (Fig. 6b). On average, the decay with sleep time is similar for down states, delta waves and type-1 slow waves (Fig. 6c), for all the regressions tested. The decay is attenuated for SWA, notably considering the percentage of sessions with a significant regression. However, the type-2 slow waves show a systematic decrease in the slope and the percentage of sessions with significant regression (Fig. 6c).

**Figure 6-.**
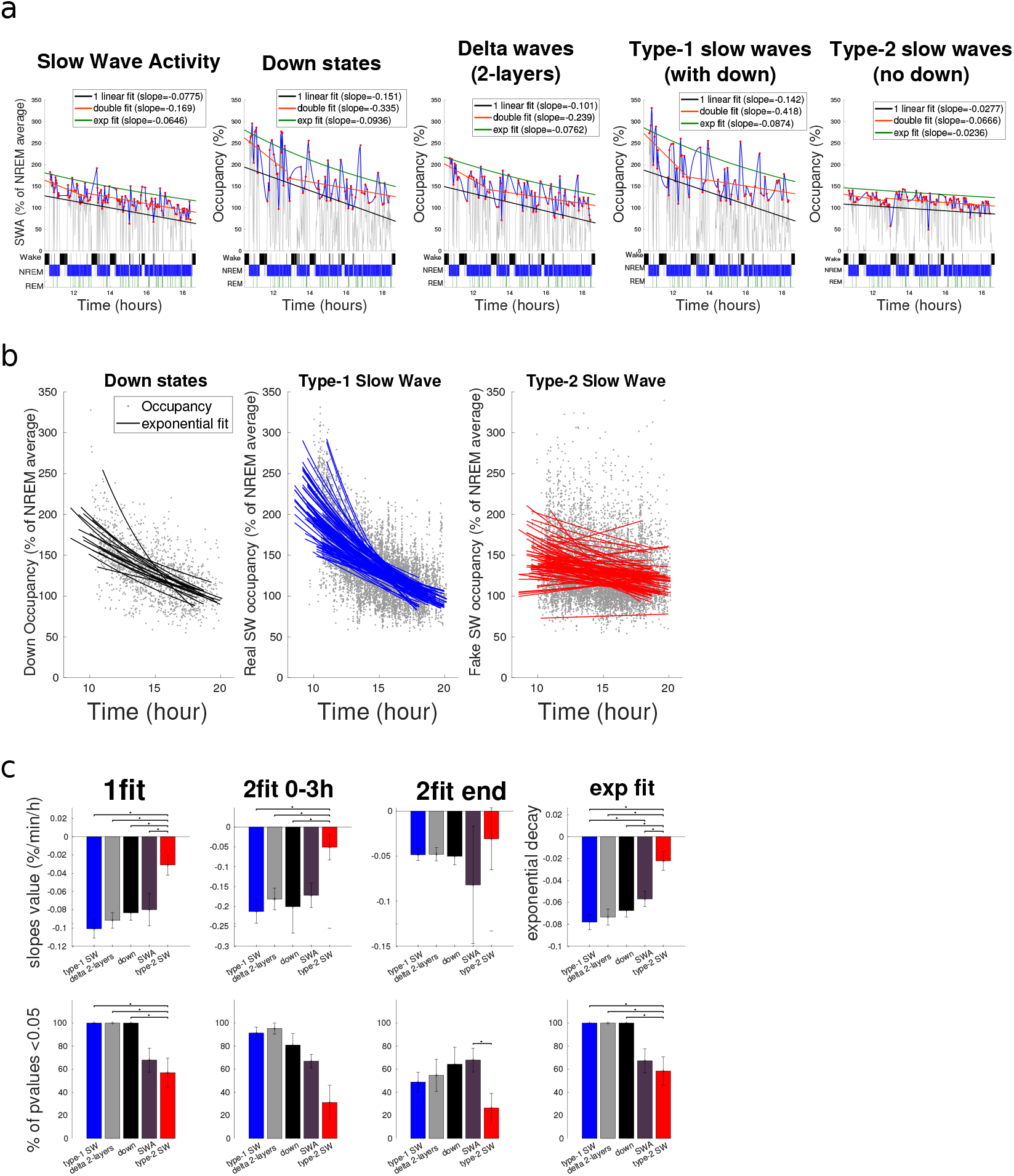
Accurate detection is crucial for the assessment of sleep homeostasis. **a**. Example of sleep homeostasis measurements for one sleep session. *Left to Right*: Slow-wave activity SWA: 0.5-4 Hz spectral power, down states occupancy (% of time in down state), delta waves occupancy (detected with 2-layers), type-1 SW occupancy (with down), type-2 SW occupancy (no down). All these measures are normalized by their NREM average value. For each curve, 3 types of regression are computed: 1 linear fit, double linear fit (fit on the first 3 hours and fit on the end) and exponential fit. For display reasons, linear 1-fit curves are shifted down and exponential fit curves are shifted up. Fits are done on peaks of homeostatic curves, in red. The hypnogram is shown below. **b**. Aggregated plot of all peaks and exponential fits. Homeostatic decrease is obvious for down states and type-1 slow waves, but not for type-2 slow waves. **c**. (top) Quantification of homeostatic measurement: type-2 SW give significantly different assessments of homeostatic decrease (Paired Wilcoxon signed-rank test, n=7). (bottom) p-value for each fit: low values for type-2 SW indicate weaker correlations between their evolution and time (Paired Wilcoxon signed-rank test, n=7).

**Figure 7-.**
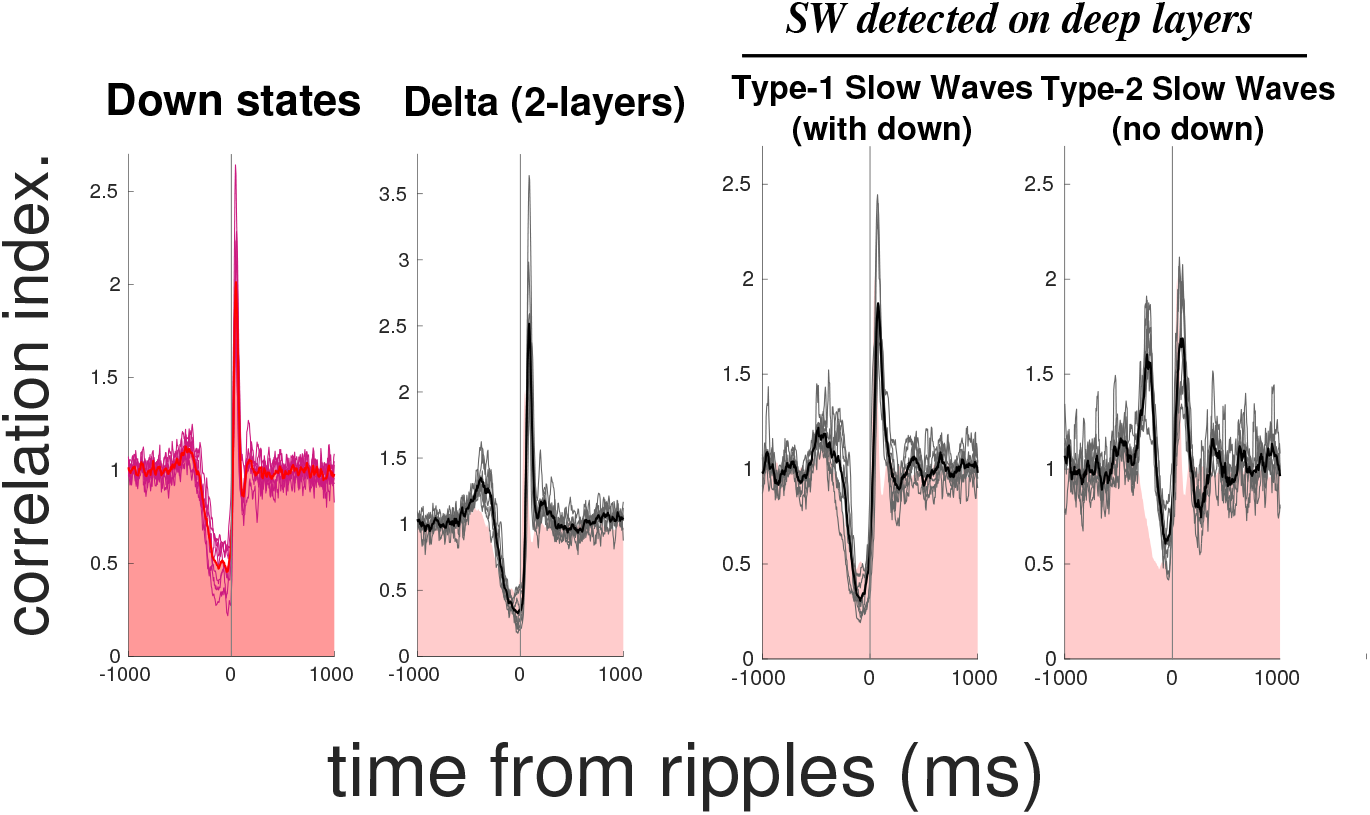
Time scale relationship with sharp-wave ripples. Cross-correlogram between ripples and down states / delta / slow waves (ripples at t=0ms, n=6). *From left to right*: down states, delta waves from 2-layer method, type-1 slow wave (deep), type-2 slow wave (deep). Red areas correspond to down state cross-correlogram.

### Interaction between hippocampal SWRs and the slow waves

There are now strong pieces of evidence showing that the occurrence of hippocampal SWR and delta waves or down states are not independent. However, some studies found that SWRs occur before the up states (thus at the end of the down states), some found that delta waves followed SWRs and others that delta waves occur just before and after SWRs [14]–[16]. We believe that these discrepancies can be explained in part by the confusion between slow waves and delta waves. We therefore quantified the cross-correlogram between hippocampal SWRs and cortical events: down states, delta waves and the two types of slow waves. Both the down states and the delta waves, we observed that they followed SWRs with a strong and narrow peak at 45ms (for the beginning of the down states) and 85ms for the peak of the delta waves. This suggests that the cortical excitation induced by the hippocampal SWRs could induce a delta wave. We noticed the same relation between SWRs and type-1 slow waves. On the contrary, for the type-2 slow waves, we observed a peak before and after the hippocampal SWRs, suggesting that the peak observed before SWRs in the literature might be due to type-2 slow waves, due to the inappropriate identification of true delta waves.

Type 2 slow waves are thus electrophysiological events that may have an important role for memory consolidation but have no relationship with down states and do not show a homeostatic regulation. Altogether, our study shows that delta waves are a subpopulation of slow waves and are specifically regulated in a homeostatic way.

## Discussion

Cortical slow waves and delta waves are hallmarks of NREM sleep and key elements in physiological processes such as memory consolidation and sleep homeostasis [16], [31]. However, the terminology used to dissociate between slow electrophysiological events during NREM sleep is still rather confusing: slow oscillation, delta wave, K-complex, delta oscillation… It is, for instance, difficult to differentiate K-complexes and slow oscillations. For the latter dissociation, it is generally accepted that K-complexes are found isolated, but waves of approximatively the same shape will be called slow oscillation when occurring in bursts [32]. However, there is no clear evidence of whether the neuronal activity associated with these LFP/EEG events is different or not. Similarly, the distinction between K-complexes and delta waves is rather vague. Both are supposed to be associated with a cortical down state, but there is still a controversy regarding the existence of an increase in excitation at the end of the event, which would be a characteristic of the K-complex [33]–[35]. Finally, slow oscillations are also supposed to be related to up and down states fluctuations [27], but it is sometimes difficult to differentiate the contribution of the delta waves in this process. Moreover, the term oscillation made people use the phase to define the up and down states. However, down states are unitary events that can occur sometimes in a regular fashion, but the use of a phase to identify their occurrence is sometimes misleading (see for instance [36]).

One conservative hypothesis is that a delta wave is the neuronal correlates of a down states, even if it is still difficult to understand how a cessation of neuronal activity is able to generate an electric dipole in the cortex (but see [30]). The duration of both delta waves and down state is of the order of 200-400ms [33], [37]. The occurrence of delta waves during sleep will thus enhance the 2-4Hz (delta) frequency range. Importantly, the occurrence of the delta wave is not random but follows a certain kind of rhythmicity. The slow oscillation would thus correspond to the underlying process that controls the occurrence of delta waves [24]. During sleep, the Up states are longer that the down states leading to a lower frequency of the order of 1Hz [37].

According to this conceptual framework, the precise identification of delta waves is thus crucial. Yet, the NREM sleep is full of large fluctuations of slow frequency that can easily be detected as a delta wave.

In this study, we introduce a new distinction between slow waves, i.e. large amplitude waves on filtered LFP cortical signals, and delta waves, characterized by an inversion between deep and superficial layers; and we separate type-1 slow waves, that co-occur with a down state, and type-2 slow waves that are not associated with a down state.

Detection of slow waves as large amplitude waves during NREM sleep reveals that a large portion of them do not share the properties of delta waves. We show that 1) only true delta waves and type-1 slow waves undergo strong homeostatic regulations and 2) type-2 slow waves create confusion about the temporal link with hippocampal sharp-wave ripples.

For instance, in one of the first articles that investigated the link between delta waves and ripples, Sirota and colleagues detected delta waves as positive deflections in the deep layer above 3 SD. They observe temporal coordination, with the ripples coming preferentially before the delta waves [14]. On the contrary, another study found the exact opposite with delta waves before the hippocampal SWRs [38], and some study found the delta waves before and after the SWRs [16], [18]. According to our results, the putative delta waves found before the SWRs would correspond to type-2 slow waves with no down state associated. This does not mean that type-2 slow waves are not relevant electrophysiological events. It only means that they are not related to a down state and that the waves found before and after the SWRs are not similar.

The same problem of interpretation is likely to occur in the recent article that tried to make a dissociation between slow oscillation and delta waves [36]. In this article, the difference is made according to the shape of the wave, and more precisely on a bump before the wave. When looking at the raw data presented, the waves are of small amplitude, that would correspond in our study to group 1-3, meaning that the error of identification of delta waves is high. This is confirmed by the analysis of spiking activity that shows only a moderate decrease in firing rate at the time of the detected event (see their figure 2e). This makes the interpretation of their results extremely difficult since, according to our current study, almost half of the events detected do not correspond to down states. The manipulation with optogenetic tools could have an effect either through the manipulation of down states or through the manipulation of type-2 slow waves that are associated with the SWRs.

Similarly, most of the studies on sleep homeostasis focus on the slow waves as a marker of sleep pressure. Yet, they address this issue using the spectral power within the 0.5-4Hz frequency band [20], [39], or the extrema of the large amplitude and low frequency fluctuations during NREM sleep [40]. Both methods will detect the two types of slow waves. This is a minor issue if the purpose is to quantify the decay with sleeping time since we observed this effect even when adding noise related to the type-2 slow waves. However, this becomes an issue when we try to understand the neurophysiological correlates of this decay and if we want to investigate whether the slow waves are more than a marker of sleep pressure. We need to understand precisely what is modulated by sleep pressure to understand how this process could be involved in the restorative function of sleep. According to our study, this would be linked to the process involved in the UP and DOWN state fluctuations.

Altogether, our results confirm the importance of a robust and accurate definition of delta waves in order to understand the fine neurophysiological mechanisms at stake during NREM sleep. We propose to use systematically the difference between the superficial and deep layers to identify delta waves in a reliable manner.

## Materials and Methods

### Experimental Design

Controlled laboratory experiments were conducted on a total of 9 C57Bl6 male mice (*Mus musculus*), 3–6 months old. No blind experiment or subject randomization was used.

### Surgical protocols and behavior experiments on mice

All mice underwent stereotaxic microsurgery for electrode implantation. Simple tungsten wires were lowered in the prelimbic cortex (AP +1.9, ML 0.4, DV −1.6), in the CA1 pyramidal layer of hippocampus (AP +2.2, ML +2.0, DV −1.0) and the deep layer of the olfactory bulb (AP +4.70, ML -0.6, DV -0.9) of the right hemisphere. All mice were implanted with tetrodes in the prelimbic cortex to record single-unit activity. During recovery from surgery (>7 days) and during all experiments, mice were housed in an animal facility (12h light / 12h dark, constant light and monitored temperature), one per cage, and received food and water ad libitum. All mice were free of any manipulation before being included in this study. Natural sleep in mice home cage was continuously recorded (10am to 8pm).

All behavioral experiments were performed in accordance with the official European guidelines for the care and use of laboratory animals (86/609/EEC) and in accordance with the Policies of the French Committee of Ethics (Decrees n° 87–848 and n° 2001–464). Animal housing facility of the laboratory where experiments were made is fully accredited by the French Direction of Veterinary Services (B-75-05-24, 18 May 2010). Animal surgeries and experimentations were authorized by the French Direction of Veterinary Services for K.B. (14-43).

### Electrophysiological recordings and analysis

Signal was sampled at 20 kHz, digitalized and amplified by INTAN system (RHD2000-series). Local field potentials (LFP) were sampled and stored at 1.25 kHz. Recordings were band-pass filtered between 0.6-9kHz, processed using NeuroScope [41] and single-units were sorted using KlustaKwik [42].

### Sleep scoring

Sleep and wake episodes were distinguished by automatic k-means clustering of the gamma (50-70 Hz) distribution extracted from the power spectrograms of olfactory bulb LFP signal from the whole recording session. Sleep corresponds to low power of gamma, wake to the rest of the recording session [43]. REM and NREM sleep were distinguished by automatic k-means clustering of the theta/delta ratio extracted from the power spectrograms of hippocampal LFP signal during the episodes where the animal was asleep. REM sleep corresponds to high theta/delta ratio periods, NREM sleep to the rest of sleep.

### Down state detection

Offline down states detection used multi-unit activity (MUA) computed from the neuronal population of the prelimbic cortex. MUA was computed as the sum all spikes in time windows of 10ms. First, periods of MUA equal to zero of at least 30ms were detected and merged if spaced by 10ms. Then, periods that last more than 75ms occurring in NREM sleep were classified as down states.

### Slow wave detection

Offline slow wave detection used one LFP channel recorded in the prefrontal cortex. Channels were classified as deep or superficial in function of their responses on down states: deep for a positive response and superficial for a negative response. For deep/superficial channels, periods during which the filtered LFP signal (in the 1-5Hz frequency band) was above/below a threshold of 2 standard deviations (SD) were selected. A threshold of 1 SD is used to determine the starts and ends of these periods. Then, periods that last more than 75ms occurring in NREM sleep were classified as slow waves.

### Delta wave detection with the 2-layer method

Offline delta wave detection used two LFP channels recorded in the prefrontal cortex, ideally one in a superficial layer and the other in a deep layer. In the absence of the latter configuration, two channels at different depths were selected, so that the dipole generated by delta waves was clearly apparent. Those two signals were band-pass filtered (1-20Hz) and subtracted to obtain the differential signal between deep and superficial layers. Delta waves were then defined as extrema of this filtered differential signal in the 1-12Hz frequency band, above a threshold of 2 standard deviations (SD) defined during sleep, and that last at least 75ms. A threshold of 1 SD is used to determine the starts and ends of delta waves.

### False positive quantification and co-occurrence with down states

In figure 4, delta waves or slow waves were defined as false positive when they did not co-occur with a down state, which means that the periods do not overlap. Superficial slow waves were given a margin of 50ms for overlapping with down states, as superficial layers responses to slow waves occur later than deep layers. False positive rate was measured as the percentage of slow/delta waves not associated with a down state.

### Homeostasis analysis and slow wave activity

*S*low wave activity (SWA) was defined as the power in the slow wave range (0.5-4Hz). Values were subsequently normalized to the mean over all NREM sleep epochs of the recording.

### Homeostasis analysis and occupancy analysis

The occupancy is calculated as the percentage of epoch duration in time windows of 60 seconds. It can be computed for down states, slow waves and delta waves. Its evolution through the recording session is then assessed for further homeostasis analysis.

### Homeostasis analysis and linear/exponential regressions

Regressions are computed to estimate homeostasis decreases of SWA or NREM occupancy of down states, slow waves or delta waves. Regressions are performed on NREM local maxima of SWA/occupancy evolution. 3 types of regressions are used :

*One fit:* linear regression on the whole sleep session

*Two fits* (double linear regression): regression on the first 3 hours of sleep session + regression on the rest of the recording.

*Exponential fit:* exponential fit on the whole sleep session, in the form *a*. exp (*b. t*)

Slopes are compared for linear regressions, and exponential decays *b* are compared for exponential fit.

## Author contributions

K.E did the experiments and analyzed the data. G.L, S.B and M.L did several experiments included in the dataset. K.E and K.B wrote the manuscript with the help of S.B.

